# Atomic-level characterization of the conformational transition pathways in SARS-CoV-1 and SARS-CoV-2 spike proteins

**DOI:** 10.1101/2022.11.29.518406

**Authors:** Dylan S Ogden, Mahmoud Moradi

**Affiliations:** Department of Chemistry and Biochemistry, University of Arkansas, Fayetteville, Arkansas 72701, U.S.A.

## Abstract

Severe acute respiratory syndrome (SARS) coronaviruses 1 and 2 (SARS-CoV-1 and SARS-CoV-2) derive transmissibility from spike protein activation in the receptor binding domain (RBD) and binding to the host cell angiotensin converting enzyme 2 (ACE2). However, the mechanistic details that describe the large-scale conformational changes associated with spike protein activation or deactivation are still somewhat unknown. Here, we have employed an extensive set of nonequilibrium all-atom molecular dynamics (MD) simulations, utilizing a novel protocol, for the SARS-CoV-1 (CoV-1) and SARS-CoV-2 (CoV-2) prefusion spike proteins in order to characterize the conformational pathways associated with the active-to-inactive transition. Our results indicate that both CoV-1 and CoV-2 spike proteins undergo conformational transitions along pathways unique to each protein. We have identified a number of key residues that form various inter-domain saltbridges, suggesting a multi-stage conformational change along the pathways. We have also constructed the free energy profiles along the transition pathways for both CoV-1 and CoV-2 spike proteins. The CoV-2 spike protein must overcome larger free energy barriers to undergo conformational changes towards protein activation or deactivation, when compared to CoV-1.

## Introduction

Severe acute respiratory syndrome coronavirus 2 (SARS-CoV-2) mediates viral entry into host cell receptors ^1–4^ and is the major cause for the coronavirus disease pandemic in 2019 (COVID-19) ^5–9^. SARS-CoV-2 is by far the most effective coronavirus, among SARS coronavirus 1 (SARS-CoV-1) and Middle East respiratory syndrome (MERS) ^10^, in terms of being easily transmissible to millions of people around the world ^11^. Genomic analysis of SARS-related coronaviruses has indicated the coronavirus spike protein is the most variable region of the entire genome ^5^. This particular protein is involved in host cell recognition, by binding the human angiotensin-converting enzyme 2 (ACE2) receptor, and viral entry by mediating membrane fusion. Interestingly, the CoV-2 spike protein has been shown to have higher affinity for human ACE2 than the CoV-1 spike protein ^10^.

SARS-CoV-2 must undergo large-scale confromational changes in the receptor binding domain (RBD) in order to interact and bind with ACE2 ^1–4^. The RBD transitions from the inactive state (“down”) to the active state (“up”) to facilitate binding of the receptor binding motif (RBM) with ACE2 ^6^. Recent Cryo-EM studies have successfully resolved structures of both spike proteins in the inactive state, active unbound state, and active ACE2-bound state ^10, 12–15^. However, cryoEM and X-ray crystallography studies capture static pictures of specific protein conformations and do not provide detailed information on the dynamic behavior associated with the major protein conformational changes ^16–18^. In addition, given the substantial differences in the experimental and physiological conditions, it is not clear whether all relevant conformational states are captured using techniques such as cryo-EM. For instance, a recent single-molecule fluorescence resonance energy transfer (smFRET) study has captured an alternative inactive conformation for the CoV-2 spike protein ^19^ that is not consistent with those obtained from cryo-EM. It is important to decipher the differential conformational landscapes of the CoV-1 and CoV-2 spike proteins in terms of both important functional states and their dynamics. Understanding the activation mechanism of the coronavirus spike proteins and its potential variant-dependent nature is key to the development of efficient coronavirus vaccines and therapeutics with long-term efficacy.

All-atom molecular dynamics (MD) simulations can provide dynamic pictures of biomolecular processes at a detailed molecular level. Often times MD simulations are often unable functionally relevant events, such as those involved in spike protein activation, tend to extend beyond the the time scales of most MD simulations. The scope of MD simulation studies of large protein complexes is most often limited to the local conformational changes of individual states starting from available crystal structures or homology models ^20–30^. An efficient computational framework for describing large-scale conformational changes using a combination of several distinct enhanced sampling techniques, without compromising atomic details, has been previously developed ^31–34^ and employed in this work.

In order to generate functionally relevant state transitions between protein activation and inactivation in both SARS-CoV-1 and SARS-CoV-2, we have employed Steered Molecular Dynamics (SMD) to generate initial transition pathways and followed by String Method with Swarms of Trajectories (SMwST) for further transition path characterization and refinement in high-dimensional collective variable spaces. Finally, Bias Exchange Umbrella Sampling (BEUS) is employed to effectively sample along the transition pathways using optimized reaction coordinates in order to determine the free energy associated with the conformational changes.

By employing a novel enhanced sampling scheme, we have conducted an extensive computational study to characterize the transition pathways for both SARS-CoV-1 and CoV-2 spike proteins. Our results provide details indicating two very different transition pathways associated with the deactivation of the active protomer in both systems. We also observe multiple local interactions along the pathways in which the active RBD of each protein forms key salt-bridges along the pathways with an adjacent protomer’s (promters B and C) RBD, N-terminal domain (NTD), and S2 domains, suggesting the forming and breaking of these salt-bridges may be required in the activation/inactivation process and provide insights into complex multi-step state transitions specific to CoV-1 and CoV-2. Furthermore, we observe differing dynamics of the inactive protomer RBDs for both CoV-1 and 2 along the transition pathways. Finally, the determined free energy profiles for both proteins have been constructed and further our results from our previous study ^35^.. Protein activation or deactivation in CoV-2 is hindered by a higher free energy barriers than CoV-1.

## Methods

### All-atom equilibrium MD simulations

We have used all-atom equilibrium MD simulations to characterize the active-to-inactive conformational transition pathways of the two spike proteins SARS-CoV-2 and SARS-CoV-1. Our simulations were derived from the two cryo-EM structures of the SARS-CoV-2 spike protein (PDB entry:6VYB) ^10^ and SARS-CoV-1 spike protein (PDB entry:5×5B) ^13^, both containing one active promoter. The protein was solvated in a box of TIP3P waters, and 0.15 M NaCl (in addition to the counterions used to neutralize the protein) using CHARMM-GUI ^36, 37^. The box size for the CoV-2 active model was 198 × 198 × 198 Å ^3^ with 850000 atoms. The box size for the CoV-1 active model was 193 × 193 × 193 Å ^3^ with 750000 atoms.

All simulations were conducted using the NAMD 2.13/14 ^38^ simulation package with systems parameterized with the CHARMM36m all-atom additive force field ^39^. As described in previous work ^35^, each system was energy-minimized for 10,000 steps using the conjugate gradient algorithm ^40^. Both systems were relaxed using restrained MD simulations in a step-wise manner using the standard CHARMM-GUI protocol ^36, 37^ (“relaxation step”). Backbone and sidechain restraints were used for 10 ns with a force constant of 1 kcal/mol.Å ^2^ and 0.5 kcal/mol.Å ^2^ respectively (“restraining step”). The systems were then equilibrated with no bias for another 10 ns (“equilibration step”). The initial relaxation was performed in an NVT ensemble while the rest of the simulations were performed in an NPT ensemble. Simulations were carried out using a 2-fs time step at 310 K using a Langevin integrator with a damping coefficient of *γ* = 0.5 ps^−^1. The pressure was maintained at 1 atm using the Nose-Hoover Langevin piston method ^40, 41^. The smoothed cutoff distance for non-bonded interactions was set at 10 to 12 Å and long-range electrostatic interactions were computed with the particle mesh Ewald (PME) method ^42^.

### Steered Molecular Dynamics (SMD)

To induce activation/inactivation of a protomer initially in the inactive/active conformation, we defined collective variables based on the the C*α* RMSD of each protomer for both the CoV-1 and CoV-2 systems. Reference coordinates were taken from the corresponding active/inactive structure for both CoV-1 and CoV-2 protomers. The atoms chosen were based on the total number of resolved and modeled residues in the CoV-2 structures. Structural analysis of CoV-1 and CoV-2 was employed to ensure that equivalent C*α* atoms were steered in all simulation sets. 1037 atoms were steered for any given protomer and the following residue range was used: 27 to 239, 244 to 315, 322 to 662, 673 to 809, and 831 to 1104. These atoms span the entire protomer, starting from the NTD and ending approximately at the C-terminus of the S2 region. A force constant of 250 *kcal/mol/*Å ^2^ was used for SMD simulations involving a single protomer. Altogether 40 simulations were conducted (2 directions x 2 proteins x 10 replicas) for 100 ns (400 ns total) of SMD. One pathway was chosen for each protein whose resulting pathway required the least amount of nonequilibrum work (active-to-inactive transition pathway) ^35^. Each of the two pathways were reduced to 100 equidistant images (conformations or windows) to represent the initial transition pathway.

### String Method with Swarms of Trajectories (SMwST)

The resulting generated pathways are far from equilibrium and require path optimization. Path refinement through the use of an automatic path finding algorithm known as string method with swarms of trajectories (SMwST) ^33^ was employed. SMwST has proven successful in generating reliable transition pathways in the past ^33, 34, 43^, while taking into account the nonlinearity of multidimensional collective variables. For the first set of SMwST, we employed 200 iterations for SARS-CoV-2 and 125 iterations for SARS-CoV-1 using the atomic coordinates of the C*α*, with a force constant of 10 kcal/mol.Å ^2^, atoms of the active protomer as the collective variable to characterize the transition pathway from active to inactive, which encompassed 1037 atoms in both systems, the same as SMD. Both systems utilized 2000 replicas in total (i.e. 100 images x 20 copies/image), each iteration of SMwST included 10 ps of restraining followed by 1 ps of restraint release. The total simulation times were 4.4 *μ*s for SARS-CoV-2 and 2.75 *μ*s for SARS-CoV-1.

A second set of SMwST was used to further characterize the transition pathways using collective variables to more accurately describe the conformational changes. In our approach, we employed more system-specific collective variables such as the “orientation quaternion” ^31, 44–46^. Orientation quaternions have been proven to be efficient collective variables at inducing a rotational change of a given molecular domain or can be utilized more simply by restraining a domain’s orientation. For each system we employed four collective variables, the orientation quaternion of the three domains: RBD, NTD, and S2. Additionally, the distance (distance vector) between the RBD and S2 domain (active protomer only) was used as the fourth collective variable. The force constants used were 200 kcal/mol.Å ^2^ for distance and 500,000 kcal/mol.rad^2^ for the orientation quaternion collective variables. Furthermore, 200 iterations of SMwST were employed for CoV-2 and 120 iterations for CoV-1. Each iteration of SMwST included 10 ps of restraining and 2 ps of restraint release. Similar to the first set, each system contained 2000 replicas (i.e. 100 images x 20 copies/image). The final simulation times were 4.4 *μ*s for CoV-2 and 2.88 *μ*s for CoV-1. In total the simulation time for CoV-2 is 8.8 *μ*s and 5.63 *μ*s for CoV-1, and in grand total: 14.43 *μ*s for all SMwST simulations.

### Bias Exchange Umbrella Sampling (BEUS)

Once optimized pathways have been generated from SMwST, the pathways were be further characterized by calculating the free energy along the transition paths. The novel Bias Exchange Umbrella Sampling (BEUS) scheme utilized in this work, allows for very accurate sampling along a transition pathway best represented by the optimal transition pathways generated from the SMwST simulations ^31–34^. The two converged SMwST simulations were reduced to 100 replicas, 1 replica extracted from each of the 100 images. We performed 25 ns of BEUS, a replica exchange was attempted every 2 ps, in the reparameterized collective variable space described above, to provide enough sampled dynamics for accurate construction of active-to-inactive free energy profiles. The force constants used for each of the collective variables was 2000 kcal/mol.rad^2^ for the orientation quaternion restraints (both systems) and for the distance restraints 2 kcal/mol.Å ^2^ for CoV-2 and 0.5 kcal/mol.Å ^2^ for CoV-1. The exchange rates for 100 replicas in both CoV-1 and CoV-2 were as follows: 64% for CoV-1 and 63% for CoV-2.

### Theoretical Framework

The SMwST algorithm ^33, 47^ starts from an initial string, defined by N points/images {**x**_*i*_}, where *i* is any integer from 1 to *N*. Colvar *ζ* primarily defines the biasing potential, which is 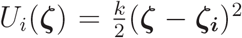 for *M* copies of image *i*. The initial values for the image centers are determined from the initial string: *ζ*_***i***_ = *ζ***(x**_***i***_). The SMwST algorithm consists of three iterative steps as follows. (Step I) Restraining: Each system is restrained for *τ*_*R*_ (restraining time) using the harmonic potential described above centered at the current image *ζ*_***i***_. (Step II) Drifting: The simulations are continued after being released from restraints for *τ*_*D*_ (drifiting time). (Step III) Reparametrization: The new center for each image *i* is determined by averaging over all observed *ζ*(**x**) values of M systems associated with image *i* at time *τ*_*R*_ + *τ*_*D*_ and using a linear interpolation algorithm to keep the image centers equidistant. By iterating over these steps, the string will converge to the zero-drift path, around which the string centers oscillate (upon convergence). The zero-drift path is an approximation of the MFEP ^48, 49^.

Once the MFEP (parametrized by *ξ*) is known, *F* (*ξ*) can be estimated using a generalization of US ^50^, termed BEUS ^33^. Similar to the SMwST method, *ξ* is discretized and *N* umbrella windows/images are defined with biasing potentials 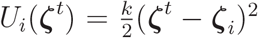 for *i* = 1,…,*N*. This scheme can be thought of as a 1D US along the reaction coordinate *ξ* with an additional restraint on the (shortest) distance from the *ζ*(*ξ*) curve. Perturbed free energies *F*_*i*_ = *F* (*ζ*(*i*)) can be estimated (up to an additive constant) by self-consistently solving the equations ^51–53^:

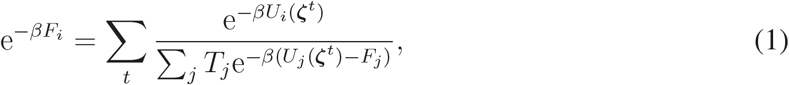

in which −_*t*_ sums over all collected samples (irrespective of which replica or image they belong to) and *T*_*j*_ is the number of samples collected for image *j*.

With appropriate reweighting, PMF can be reconstructed in any arbitrary collective variable space, given sufficient sampling in that space. *w*^*t*^, the unnormalized weight of configuration **x**^*t*^ can be estimated via ^52^:

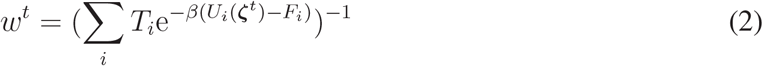

in which {*F*_*i*_} are estimated via Equation (1). Alternatively ^52^, one may estimate {*w*^*t*^} and {*F*_*i*_} by iteratively solving Equation (2) and:

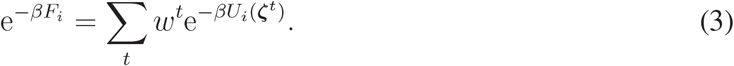

The PMF in terms of ***η***(**x**), an arbitrary collective variable, is estimated (up to an additive constant) as:

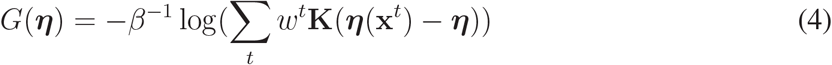

in which **K** is a kernel function. The above estimator is not accurate if the sampling in ***η***(**x**) is not converged which is the case if ***η***(**x**) has a slow dynamics and is not strongly correlated with *ζ*. For the special case of ***η*** = *ζ*, the perturbed free energies {*F*_*i*_} can be used directly to estimate the PMF only within the stiff-spring approximation.

Finally, for averaging an arbitrary quantity *A*(**x**) along the pathway *ζ*(*ξ*), one may use the weighted average *Ā*(*ξ*) = −_*t*_ *w*^*t*^*A*(**x**^*t*^)*δ*(*ζ*^*t*^ − *ζ*(*ξ*)). However, the unweighted estimator *Ā*_*i*_ = \*A*(**x**^*t*^)1_*i*_ is often more efficient. 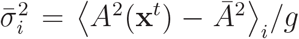 provides an estimate for the variance, given 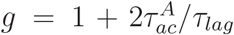 is the statistical inefficiency in which 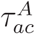 is the autocorrelation time associated with quantity *A* and *τ*_*lag*_ is the lag time between the data points used in the analysis ^32^.

## Results

We recently performed a combination of all-atom microsecond-level and SMD simulations of the non-glycosylated spike proteins SARS-CoV-1 and the original SARS-CoV-2 strain ^35^. In all but one of the systems simulated, no large-scale conformational change occurred within the timescales of the unbiased simulations. The simulation that was initiated from the active conformation for CoV-1 transitioned from active to a “pseudo-inactive” and provided details on a number local interactions between the CoV-1 spike NTD and RBD, some of which are unique to CoV-1, and were directly involved in the deactivation of the spike protein. Furthermore, the SMD simulations showed that it is relatively difficult for the CoV-2 spike protein to undergo a large-scale conformational transition between active and inactive states, when compared to the CoV-1 spike protein.

To further the work previously employed, we aimed to more accurately characterize the conformational transition of the active protomer in both CoV-1 and CoV-2 using a computational protocol that has proven to be reliable at accurately describing large-scale conformational changes ^31–34^. Using the pathways that were generated from the previous sets of SMD simulations ^35^, we chose one pathway for both CoV-1 and CoV-2 to be used as the initial defining pathway to be further characterized by the SMwST simulations. These particular pathways were derived from the sets of simulations in the direction of active-to-inactive. In order to determine the appropriate pathways to be utilized, we employed a “work-based” analysis when choosing them, (i.e. the pathways that required the least amount of nonequilibrum work) under the assumption that the pathways generated were energetically favored. Currently, the pathway that connects the active and inactive states in both proteins is not known but the cryo-EM structures provide details on the two end states (active and inactive) ^10, 13^.

The work that is presented here is primarily derived from an extensive set of SMwST simulations (data reported is generated from the last iteration of the final set of string method simulations) that employed string method using different definitions for the collective variables used in biasing the active protomer. The first set focused on biasing the atomic coordinates of the of C*α* atoms that encompass the active protomer. Atomic coordinate ^43^ biasing has shown to be a useful application for SMwST simulations when characterizing transition pathways. The second set employed distance and orientation quaternion based collective variables in order to describe the conformational changes of the active protomer deactivation more appropriately. Supplementary figures 1-4 show the convergence of the two sets of string method simulations with collective variables that provide insight into the global conformational changes of the CoV-1 and CoV-2 spike protein’s active protomer along the transition pathways.

### Global protein conformational changes

With the transition pathways characterized, the global protein conformational changes along the deactivation pathways can be described. Figure 1 A-D, emphasizes the similarities and differences between two proteins when in the active and inactive conformations, with CoV-1 adopting a more “active” conformation (Fig.1A) as the RBD appears to be pointed straight up, while the RBD in CoV-2 is angled downward in the direction toward NTD_*B*_ (Fig.1C). The inactive conformations (Fig.1B,D) show a noticeable difference in the RBD orientations with the RBD in CoV-1 being slightly tilted downwards and away from the adjacent RBDs and the RBD in CoV-2 tilted slightly upwards and towards the adjacent RBDs. Figure 1E shows the coupling between same protomer domains as the conformational changes is occurring along the deactivation pathway. The RBD in CoV-1 rotates away from the NTD before any change in the RBD-S2 distance occurs, the RBD-S2 distance remains constant at ∼93 Å until the angle between the two domains approaches closer to 90° and then the trend between the two calculations becomes quite linear. Unlike CoV-1, the transition of the active protomer is coupled linearly between the two RBD-S2 distance the RBD-NTD angle. The deactivation of the RBD in both proteins follows two different paths, the RBD in CoV-1 rotates farther from the NTD before transitioning closer to the S2 domain, while the RBD deactivation in CoV-2 is essentially negatively correlated between the two definitions. Furthermore, we have also monitored the degree of the ACE2 accessibility (Supplementary Fig.3), using a definition derived by Peng et al ^54^. The relative degree of the ACE2 accessibility for both CoV-1 and CoV-2 is in line with the thresholds of 52.2°-84.8°, with ACE2 being completely inaccessible for images (windows) *>*50.

**Figure 1:**
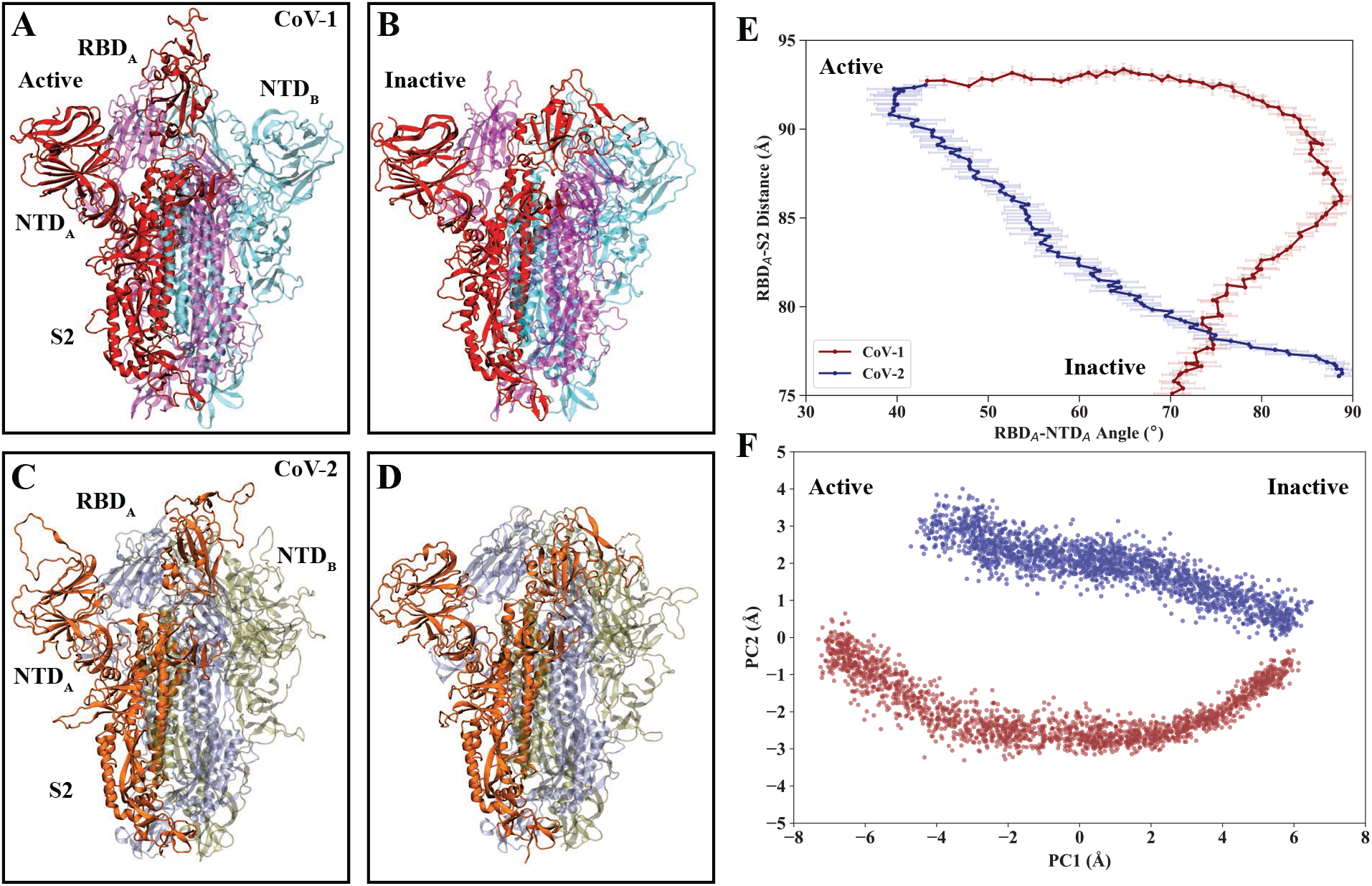
Global conformational changes of the CoV-1 and CoV-2 spike proteins. **(A, B)** Side view of active and inactive states for CoV-1. The active protomer, A, is colored in red, protomer B is colored in cyan, and protomer C is colored in magenta. **(C, D)** Side view of active and inactive states for CoV-2. Protomer A is colored in red, protomer B is colored in cyan, and protomer C is colored in magenta. Protomers B and C are represented transparent. Protomer A is colored in orange, protomer B is colored in tan, and protomer C is colored in blue. Protomers B and C (both proteins) are represented transparent. **(E)** RBD-S2 center of mass distance vs. RBD_*A*_-NTD_*A*_ interdomain angle, each domain is represented by the roll axis (longest principle axis). Each point is the average value of the 20 copies for each image and the error bars are defined by the standard deviation of the 20 copies for each image. **(F)** The first two principle components of the protomer A C*α* atoms.

To further track the conformational changes of the CoV-1 and CoV-2 spike proteins, more insight was gained by projecting the C*α* atoms of the active protomer, taken from the last frame of the SMwST simulation trajectories, onto its principle components. Figure 1F further illustrates how the two protein active protomers take two different pathways during active-to-inactive transition. PC1 and PC2 account for ∼83% of the variance with PC1 accounting for ∼63% of the variance. PC1 roughly describes the motions of the RBD rotating downward towards the inactive state, with slight deviations in both the NTD and S2 domains, and PC2 describes a rotational change of the RBD in the XY plane. Following along the pathways from active-to-inactive, both CoV-1 and 2 exhibit paths similar to those provided in Figure 1E. CoV-2 transitions linearly and CoV-1 undergoes a non-linear path in the PC space.

### Active and inactive RBD conformations

The top views of the CoV-1 and CoV-2 spike proteins (Fig.2A-F) further exhibit the differences between the two ends states (Fig.1A-D) and provide insight into two example transient conformations (image 50) along the transition pathways. The RBD, when in the active conformation, in CoV-1 is oriented equidistant between NTD_*A*_ and NTD_*B*_, while RBD_*C*_ remains in close proximity of RBD_*A*_ (Fig.2A). The active RBD in CoV-2 is positioned closer to the adjacent protomer NTD_*B*_ (Fig.2D). The transient conformation for both CoV-1 and CoV-2 spike proteins (Fig2.B,E) provides insight into the differing degrees of RBD activation halfway through the transition pathway. Further inspection of the top views (Fig.2A-F) provide insight into how the inactive RBDs (protomers B and C) adopt different orientations as the active RBD transitions towards the inactive state. All three RBDs in CoV-2 begin to pack tighter together (Fig.2E-F) as the active RBD transitions towards being inactive. In CoV-1 the RBD’s move away and position closer towards the adjacent protomer NTDs (Fig.2B,C), this is extenuated in (Fig.2C) as there is a gap between all three RBDs and much of the S2 domain is visible below (represented colorless and transparent). This is not the case for CoV-2 as all three RBDs are positioned close enough that the S2 domain is significantly less visible underneath.

**Figure 2:**
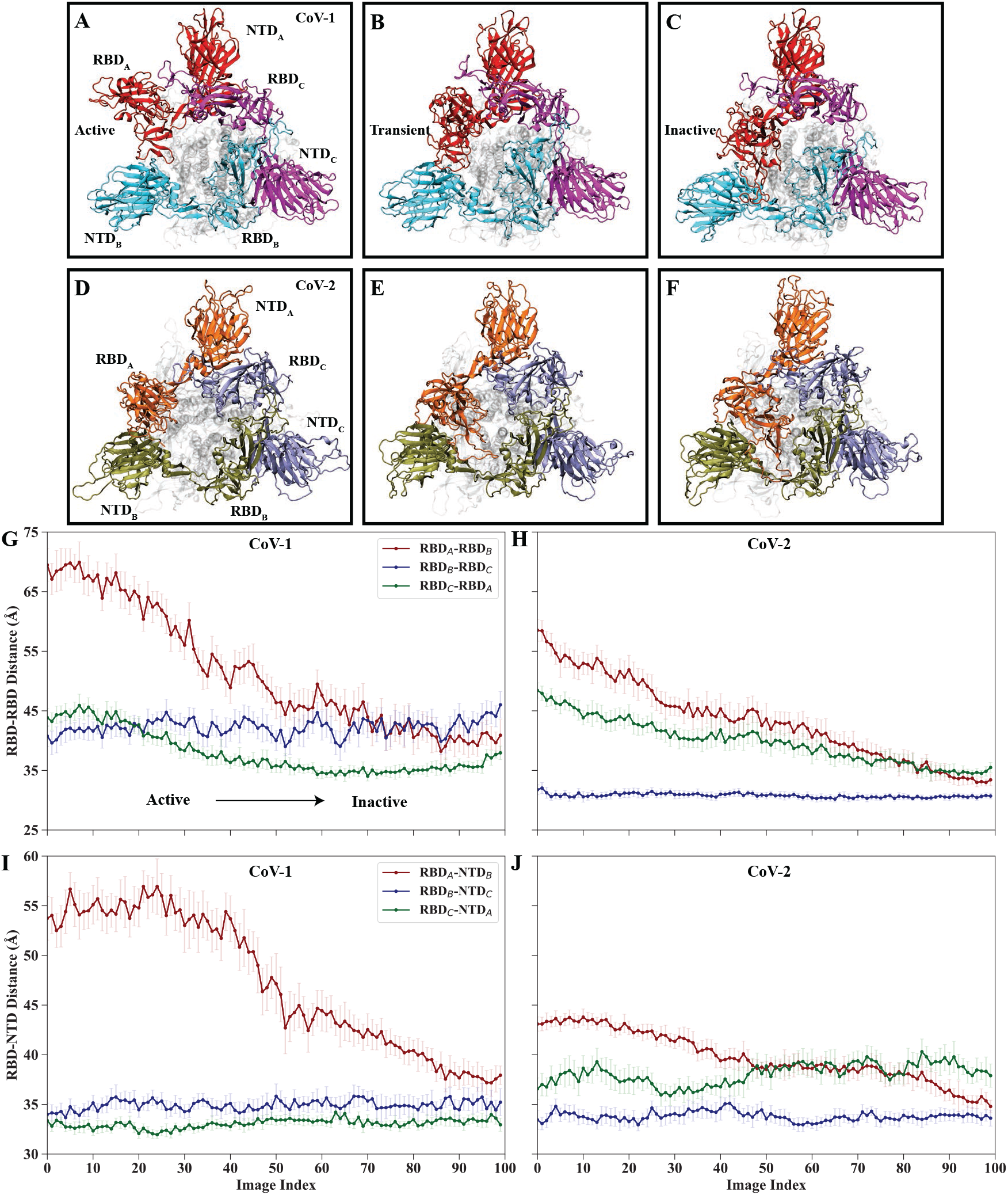
Receptor binding domains show a differential dynamic behavior. **(A-C)** Top view of active, transient, and inactive states for CoV-1. (D-F) Top view of active, transient, and inactive states for CoV-2. **(G, H)** RBD-RBD center of mass distances per image index. **(I, J)** RBD-NTD center of mass distance of each RBD and its adjacent NTD per imaged index. Each point is the average value of the 20 copies for each image and the error bars are defined by the standard deviation of the 20 copies for each image.

Figure 2 G-J tracks the inter-domain center of mass distances between the three RBD’s and the center of mass distances between each RBD and its adjacent protomer’s NTD. The trends in the RBD-RBD inter-domain distances provide further indication that the receptor binding domains tend to remain in close proximity, amongst all three, in CoV-2 (Fig.2H) and the RBDs in CoV-1 (Fig.2G) transition closer to the RBD that is adjacent or “points to” when either transitioning towards inactive or residing in the inactive conformation. Similar to the trend that is observed in Figure 1E, the RBD-NTD distance remains constant across the first 40 images and then becomes linear as it is moving closer to NTD_*B*_. The active RBD in CoV-1 moves closer to other neighboring domains (Fig.3) before transitioning closer towards NTD_*B*_. The RBDs in CoV-1 tend to adopt conformations that remain pointing towards the adjacent RBD but lie closer in the proximity to the adjacent protomer’s NTD (Fig.2A-C,I) unlike the RBDs in CoV-2, which remain in equal proximity to each of the adjacent domains (Fig.2D-F,J).

### Coupling between global and local conformational changes

Figures 3,4 illustrate the coupling between global conformational changes of the active protomer and local conformational changes through the forming and breaking of key salt-bridge interactions between the active RBD and neighboring domains along the transition pathways. These events are used to characterize the transitions of the active protomers into a number of key events along the pathways. Supplementary figures 2,3 provide the list of all key salt-bridge interactions, with respect to the image index, for both the CoV-1 and CoV-2 spike proteins.

**Figure 3:**
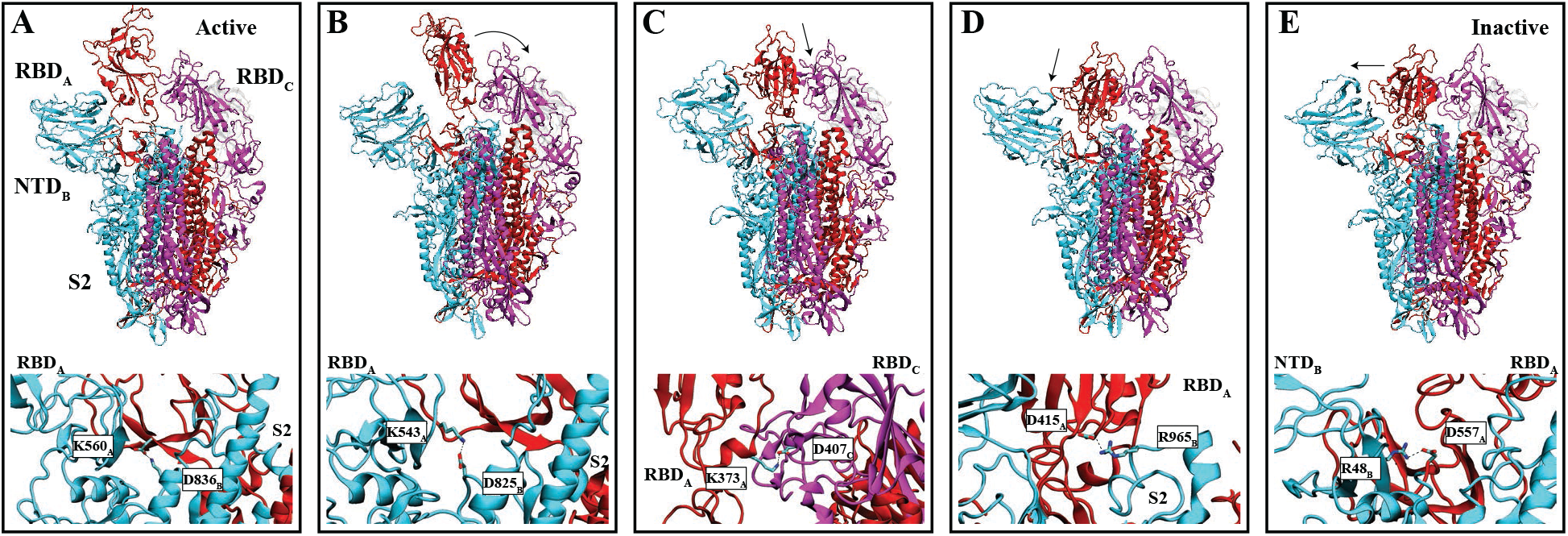
Multi-step transition pathway of CoV-1 coupled through key saltbridges. **(A-E)** Tracking the conformational changes of the CoV-1 spike protein along the transition pathway. Arrows point in the direction the active protomer RBD moves towards along the pathway. Key saltbridges are indicated below the protein snapshots that are formed along the transition pathway.

**Figure 4:**
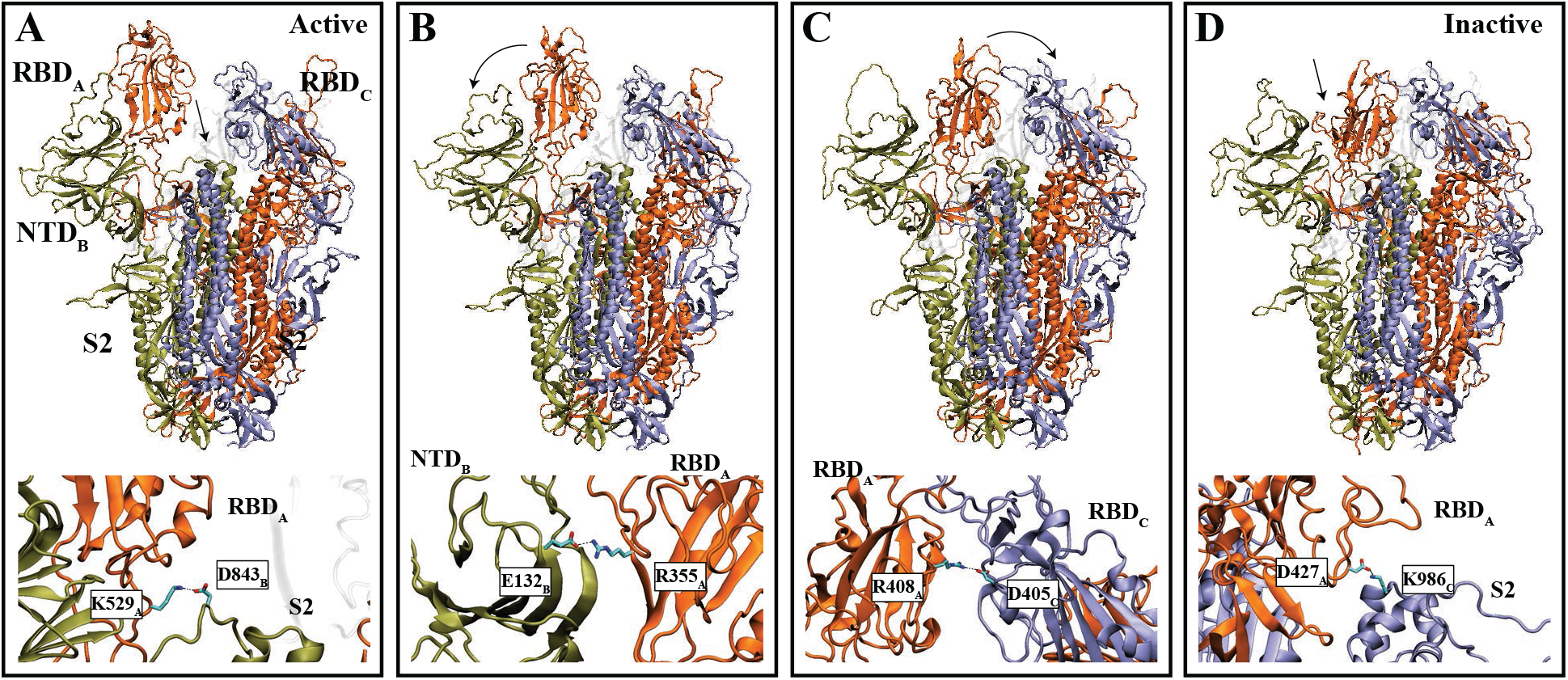
Multi-step transition pathway of CoV-2 coupled through key salt-bridges. **(A-D)** Tracking the conformational changes of the CoV-2 spike protein along the transition pathway. Arrows point in the direction the active protomer RBD moves towards along the pathway. Key salt-bridges are indicated below the protein snapshots that are formed along the transition pathway.

The RBD in CoV-1 undergoes a rotational change along its Z axis (Fig.3A) and forms a salt-bridge between D560 in the linker region between RBD_*A*_ and the S2_*A*_ and K836 of the S2 domain of protomer B. The formation of salt-bridge is quickly followed by the formation of a salt-bridge between K543 of RBD_*A*_ and D825 of S2_*B*_ (Fig.3B). RBD_*A*_ continues deactivating and transitions closer to RBD_*B*_ (Fig.3C) and forms two salt-bridges. The first being, forms very briefly and breaks, K373 of RBD_*A*_ and D407 of RBD_*C*_. The second salt-bridge forms between (Supporting Fig.2) D312 of RBD_*A*_ and K447 of RBD_*C*_. The RBD of protomer A is following by a transition downwards towards the S2 domain and transitions in the direction towards protomer B (Fig.3D,E). A salt-bridge is formed between K373 of RBD_*A*_ and E730 of S2_*B*_ as the RBD continues transitioning towards inactive. The stabilization of the inactive state is required by the formation of three salt-bridges. The first, a brief forming transient salt-bridge, between RBD_*A*_ and the NTD and S2 domains of protomer B. These salt-bridges include D572 of the linker region between RBD_*A*_ and the S2_*A*_ and R829 of S2_*B*_, D415 of RBD_*A*_ and R965 S2_*B*_, and D557 of RBD_*A*_ and R48 of NTD_*B*_.

The deactivation of RBD_*A*_ in CoV-2 begins by transitioning downwards and forms a saltbridge between K529 of RBD_*A*_ and D843 of S2_*B*_ (Fig.4A). The formation of this salt-bridge is quickly followed by the formation of a salt-bridge between K528 of RBD_*A*_ and D839 of S2_*B*_ (Supporting Fig.3). RBD_*A*_ transitions closer towards the adjacent NTD of protomer B (Fig.4 B) and forms a salt-bridge between R355 of RBD_*A*_ and E132 of NTD_*B*_. Next, RBD_*A*_ transitions towards RBD_*C*_ and forms a salt-bridge between R408 of RBD_*A*_ and D405 of RBD_*C*_. Finally, RBD_*A*_ continues transitioning towards and inactive and is stabilized by three salt-bridges, two between the RBD and NTD of protomer B, and third forming with the S2 of protomer C. The three formed salt-bridges include E484 of RBD_*A*_ and K528 of RBD_*B*_, K462 of RBD_*A*_ and D198 NTD_*B*_, and D427 of RBD_*A*_ and K986 of S2_*C*_.

### Transition path free energy calculations

We have computed the free energy profiles along the transition pathways, derived from the second set of the SMwST simulations, for both CoV-1 and CoV-2 spike proteins and utilized the same reaction coordinates described by the same collective variable space as indicated in the methods section. The free energy profiles were computed by excluding the first 5 ns, equilibrium phase, and binned the last 20 ns into 4-5ns bins for bootstrap re-sampling. The constructed free energy profiles (Fig.5) with respect to the image index (includes snapshots of the spike proteins at the associated image index, representing the minima and transition states), indicates a higher free energy barrier for Cov-2, with height 3.1 kcal.mol^−1^. The profile also contains a broad free energy well located in the region of active conformations, composed of images 5-40 and local minima with a free energy of 1.2 kcal.mol^−1^. The global minimum resides with an inactive conformation at image 90. CoV-1 also has well formed, local minima, along the active conformations, images 2-20, with a free energy of 2.0 kcal.mol^−1^. The well is followed by a lower free energy barrier that CoV-2 at 2.7 kcal.mol^−1^. The global minimum for CoV-1 is located at image 80. Interestingly, a broad free energy well is present in CoV-1, similar to CoV-2, which resides in conformations associated with being in the inactive conformation. The presence of these two very broad free energy wells are described by a higher contribution of conformational entropy, and thus, the proteins can easily occupy or transition between any one of the states that encompass these free energy wells. These results are in line with our previous work ^35^ with CoV-1 exhibiting a free energy profile that indicates a low free energy barrier when transitioning towards the inactive conformation and CoV-2 is hindered by higher free energy barriers when either transitioning towards active or inactive.

**Figure 5:**
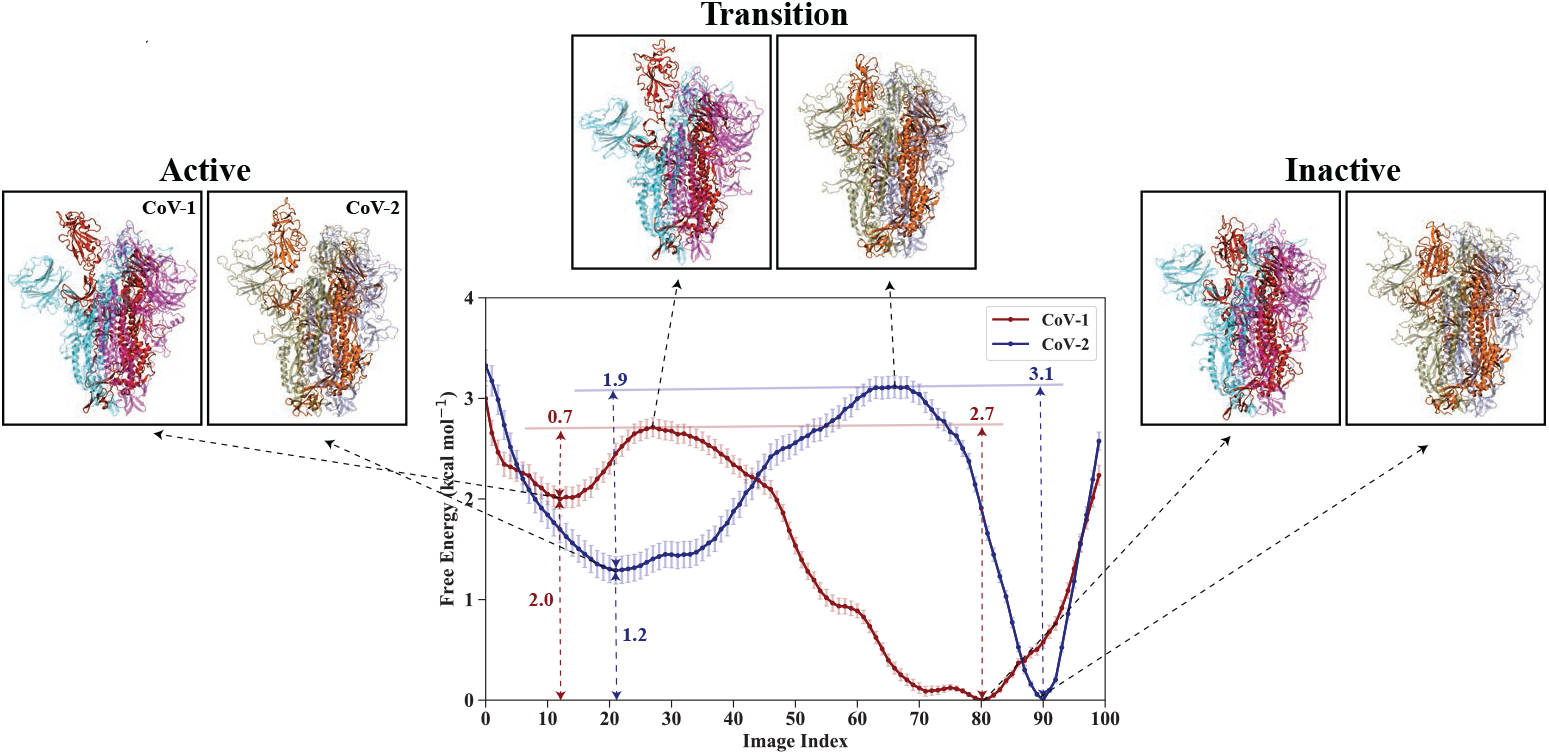
Free energy profiles for the CoV-1 and CoV-2 spike proteins. **A** Free energy profile along the transition pathways. Each image (window) represents a particular conformation for CoV-1 and CoV-2 on the discretized pathways composed of 100 images each. The error bars represent the standard deviation based on Bayesian block bootstrapping. Specific conformations of the spike proteins are utilized to represent the two minima (local and global) and the transition states. Protomers B and C are represented transparent in both proteins.

## Discussion

By employing a rigorous computational study, we have demonstrated that the SARS-CoV-1 and SARS-CoV-2 spike proteins exhibit differential dynamic behavior by characterizing the large-scale conformational changes associated with the active-to-inactive transition pathways using biased molecular dynamics simulations. More specifically, the transition pathways were first determined through the use of SMD, further refined by employing two sets of SMwST, and effectively sampled the transition pathways using BEUS to determine the associated free energy barriers along the pathways. The amount of detail provided here will allow for the testing of various experiments and even the trapping of specific intermediate states along the transition pathways.

Several experiments could be performed in order to test the results presented in our computational study. For instance, the importance certain residues play in forming key salt-bridge interactions between the active protomer RBD and adjacent domains, for both CoV-1 and CoV2, could be investigated via site-directed mutagenesis. This could shed light on some additional insights on the conformational dynamics of the CoV-1 and CoV-2 spike proteins. Additionally, smFRET experiments could be used to investigate potential, RBD-RBD, RBD-NTD, and RBD-S2 interactions by measuring the distance between fluorophores attached at specific points in each domain. Disulfide cross-linking experiments could also be used to investigate residues in the NTD, RBD, and S2 that potentially interact with each other.

MD simulations provide atomic-level elucidation of the dynamic behavior of proteins and other biomolecules ^17, 18^. Here, we have performed simulations using the non-glycosylated spike proteins of CoV-1 and CoV-2. A recent study has shown that glycosylation of the spike proteins might play an important role in the conformational dynamics of the RBD ^55, 56^. However there exists the difficulty in determining whether conformational changes occur as a result of the intrinsic protein dynamics or the differential glycosylation patterns of the CoV-1 and CoV-2 spike proteins imposed by modeling. Admittedly, it is advantageous to employ the same protocol described in this work using glycoslated models, as much is still unknown about the large-scale confromational changes that connect to the states between the active and inactive proteins.

Our results for non-glycosylated SARS-CoV-2 spike protein somewhat resembles that of the glycosylated SARS-CoV-2 spike protein, recently reported by Pang et al. ^57^ (higher and broader free energy well for the open state) but the free energy values are scaled down by a factor of ∼3 due to the absence of glycan chains ^57^. Similar reduction in the free energies was observed when the glycans were removed; however, the free energy difference between the open and closed states also disappears in the absence of the glycan chains ^57^. We believe since we have used transition path finding (string method); our results are perhaps more reliable than those obtained from direct BEUS simulations without any path optimization. Another possibility to explain the difference is that free energy values are generally collective variable dependent and we have not used the same collective variables as Pang et al. ^57^.

As discussed previously, our study primarily sheds light on the conformational dynamics of the SARS-CoV-1 and SARS-CoV-2 spike proteins. While differences in the dynamic behavior of these spike proteins almost certainly contribute to differences in transmissibility and infectivity, factors such as spike protein glycosylation and the behavior of other viral proteins also need to be considered in order to provide a more complete hypothesis. Additional experimental and computational studies are thus needed to fully investigate the differential infectivity and transmissibility of SARS-CoV-1 and SARS-CoV-2. Our simulations provide valuable insight into the dynamic behavior of the CoV-1 and CoV-2 spike proteins when transitioning between active-to-inactive. Employing the simulation protocol presented here, in the reverse direction: inactive-to-active, is required to determine the reversibility of the characterized transition pathways (i.e. global and local conformational changes and location of free energy barriers). An improved understanding of the conformational changes regarding activation or inactivation of the spike proteins, is critical to the effective development of novel therapeutics and vaccines using a structure-based drug design framework.

## Acknowledgements

Research reported in this publication was supported by the National Institute of General Medical Sciences of the National Institutes of Health under award numbers R15GM139140 and R35GM147423. This research is also supported by the National Science Foundation grant CHE 1945465 and the Arkansas Biosciences Institute. This research is part of the Frontera computing project at the Texas Advanced Computing Center, made possible by National Science Foundation award OAC-1818253. This research is also supported by the Arkansas High Performance Computing Center which is funded through multiple National Science Foundation grants and the Arkansas Economic Development Commission.

## Author Contributions

M.M. designed research; D.S.O. performed simulations and analyzed data; D.S.O. and M.M. wrote the manuscript.

